# Zika virus causes placental pyroptosis and associated adverse fetal outcomes by activating GSDME

**DOI:** 10.1101/2021.09.21.461218

**Authors:** Zikai Zhao, Qi Li, Usama Ashraf, Mengjie Yang, Wenjing Zhu, Zheng Chen, Shengbo Cao, Jing Ye

## Abstract

Zika virus (ZIKV) can be transmitted from mother to fetus during pregnancy, causing adverse fetal outcomes. Several studies have indicated that ZIKV can damage the fetal brain directly; however, whether the ZIKV-induced maternal placental injury contribute to adverse fetal outcomes are sparsely defined. Here, we discovered that ZIKV causes the pyroptosis of placental cells by activating the executor Gasdermin E (GSDME) *in vitro* and *in vivo*. Mechanistically, caspase-8 undergoes activation upon the recognition of 5’ untranslated region of viral RNA by RIG-I, followed by the stimulation of caspase-3 to ultimately escalate the GSDME cleavage. Further analyses revealed that the ablation of GSDME in ZIKV-infected pregnant mice attenuates placental pyroptosis, which consequently confers protection against adverse fetal outcomes. In conclusion, our study unveils a novel mechanism of ZIKV-induced adverse fetal outcomes via causing placental cell pyroptosis, which could be employed for developing new therapies for ZIKV-associated diseases.

**Significance statement:** Several studies have elucidated the link between ZIKV infection and congenital ZIKV syndroms (CZS), but the pathogenesis yet needs further study. Here, we reported a novel pathogenic mechanism of ZIKV which leads to pyroptosis of placental cells through activating the pyroptotic executor GSDME, rather than GSDMD. Upon ZIKV infection, GSDME-mediated pyroptosis damages the structure and function of the placenta, thereby affecting the development of the fetus and contributing to the adverse fetal outcomes. Our study highlights the importance of pyroptotic executor GSDME in regulate ZIKV pathogenicity and further confirms that placental injury caused by ZIKV infection is a key factor for CZS.

## Introduction

Zika virus (ZIKV), a mosquito-borne flavivirus, was initially identified in 1947 but received little concern until it posed a serious public health threat in the Pacific from 2007 to 2015(1). ZIKV exhibits obvious tropism and infects several immunologically privileged regions that include male and female reproductive organs, adult and fetal central/peripheral nervous system, urinary tract, and the structural and neural portions of the eye (2). Clinically, most ZIKV cases are asymptomatic or mild with low-grade fever, rash, arthralgia, myalgia, and conjunctivitis (3, 4). Nevertheless, the ZIKV infection of pregnant women still remains a major concern because the placental infection and the vertical transmission of the virus can cause adverse effects on the fetus, such as congenital ZIKV syndrome (CZS) which is characterized by microcephaly, intrauterine growth restriction, spontaneous abortion, and developmental abnormalities (5).

Although several studies have explicitly indicated a causal relationship between ZIKV infection and CZS (6-8), the underlying mechanism is not completely elucidated. On the one hand, ZIKV can infect the fetus through the transplacental route, which can lead to CZS in all trimesters of pregnancy (6, 9, 10). Studies using the brain organoids, neurospheres, and human pluripotent stem cell-derived brain cells have identified that neural stem and progenitor cells can undergo growth and developmental aberrations upon ZIKV infection, thereby resulting in microcephaly (11-13). On the other hand, the ZIKV infection-caused placental injury also likely contributes to adverse fetal outcomes. Several studies have demonstrated that ZIKV infects placental macrophages and trophoblasts, which leads to placental insufficiency and pathology (14-16). Using the mouse pregnancy models, it has been shown that ZIKV infection disrupts the architecture and function of the placenta by triggering trophoblast apoptosis and vascular endothelial cell damage (5, 8, 17, 18). Considering that placenta is infected prior to the fetus and that placenta is indispensable for the maternal-fetal nutrient exchange, it is reasonable to speculate that the placental damage during ZIKV infection could be more decisive for adverse fetal outcomes.

During ZIKV infection, the placenta can suffer from apoptosis (5, 18), which is a prototype of programmed cell death (PCD) and is crucial for reproduction, embryonic development, immunity, and viral infections. Therefore, it is meaningful to explore whether other PCD processes, such as pyroptosis, are involved in the ZIKV-induced placental damage. Pyroptosis is an innate immune mechanism against intracellular pathogens, and is featured by cell swelling and the emergence of large bubble-like structures from the plasma membrane that differentiate it from other PCDs (19, 20). Recent studies have identified gasdermin D (GSDMD) as a key pyroptosis executor, whose N- terminus domain is capable of binding to membrane lipids, phosphoinositides, and cardiolipin in order to perforate the membrane, resulting in the release of cytosolic content (21-23). Gasdermin E (GSDME), also known as DFNA5, belongs to the same superfamily as GSDMD, and was originally identified as a gene related to hearing impairment (24). GSDME is considered as a potential tumor suppressor gene in several types of cancers, such as gastric and hepatocellular carcinomas (25-27) Recently, it has also been demonstrated to mediate pyroptosis via its activated N- terminus (GSDME-N) domain that is specifically cleaved by caspase-3 and granzyme B (27, 28). GSDME can switch the caspase-3-mediated apoptosis to pyroptosis or can mediate secondary necrosis after apoptosis (29), indicating an extensive cross-talk between apoptosis and GSDME-mediated pyroptosis.

In the context of viral infections, pyroptosis is one of the imperative pathogenic mechanisms that occur during flavivirus infections. For instance, dengue virus can induce the NLRP3 inflammasome-dependent pyroptosis in human macrophages via the C-type lectin 5A, which is critical for dengue hemorrhagic fever (30, 31). Japanese encephalitis virus enhances the pyroptosis of peritoneal macrophages to raise the serum interleukin-1α level, which in turn promotes viral neuroinvasion and blood-brain-barrier disruption (32). Particularly, a recent study demonstrated the effect of ZIKV in inducing the caspase-1- and GSDMD-mediated pyroptosis of neural progenitor cells that contributes to CZS (33). In view of all these studies, it is worth exploring whether pyroptosis participates in the ZIKV-induced placental damage and adverse fetal outcomes. Here, we report that ZIKV infection activates caspase-3 via the caspase-8-mediated extrinsic apoptotic pathway, which in turn activates GSDME to induce the pyroptosis of placental trophoblasts. This phenomenon was found to associate with the ZIKV 5’UTR, but not with the ZIKV structural or nonstructural proteins, upon its recognition by the RIG-I sensor. By establishing a mouse model of ZIKV infection, we further showed that the deletion of GSDME leads to the reduction of placental damage and associated adverse fetal outcomes in infected pregnant mice. This study provides novel insights into the mechanism of ZIKV-induced placental damage and CZS; therefore, it may lead to the development of new therapeutic options to avert adverse fetal outcomes and/or miscarriages during ZIKV infection.

## Results

### ZIKV infection induces the GSDME-mediated pyroptosis in JEG-3 cells

Previous studies have shown that ZIKV infection during the first trimester of pregnancy leads to a higher chance of CZS, but the pathogenesis yet to be further studied (4, 34). Since the ZIKV-infected placenta shows obvious apoptosis of trophoblasts (5), it prompted us to hypothesize whether other routes of cell death are triggered in response to ZIKV infection. To this end, we chose a ZIKV-permissive human choriocarcinoma cell line JEG-3, which represents the features of trophoblasts and is considered a suitable in vitro model to study the first-trimester placental function (35). Upon ZIKV infection, JEG-3 cells showed evident cell swelling with balloon-like structures originating from the plasma membrane that appears to be distinct from the apoptotic blebbing (Figure 1A), displaying a characteristic pyroptotic cell morphology. Along with decreased cell viability, the swelling cells were also found to release lactate dehydrogenase (LDH, an indicator of the pyroptotic cell cytotoxicity) in an infection time-dependent manner, which depicts the rupture and leakage of the plasma membrane (Figure 1B). We next determined whether the pyroptosis executors, GSDMD and GSDME, were activated during the infection of JEG-3 cells with ZIKV. To this end, the infected cells were subjected to Western blot analysis, and the results revealed that infection led to the release of the N-terminal domain of GSDME (GSDME-N), whereas GSDMD exhibited no effect (Figure 1C), thereby implying that ZIKV infection may induce pyroptosis through GSDME. To further confirm this observation, we generated the GSDME-knockout (KO) JEG-3 cell line by employing the CRISPR/Cas9 system (Figure 1D). As expected, deletion of GSDME significantly suppressed the ZIKV-induced cell swelling (Figure 1E) together with diminished LDH release (Figure 1F). Overall, these findings demonstrate that ZIKV infection induces the pyroptosis of JEG-3 cells in the GSDME-mediated manner.

**Figure 1.**
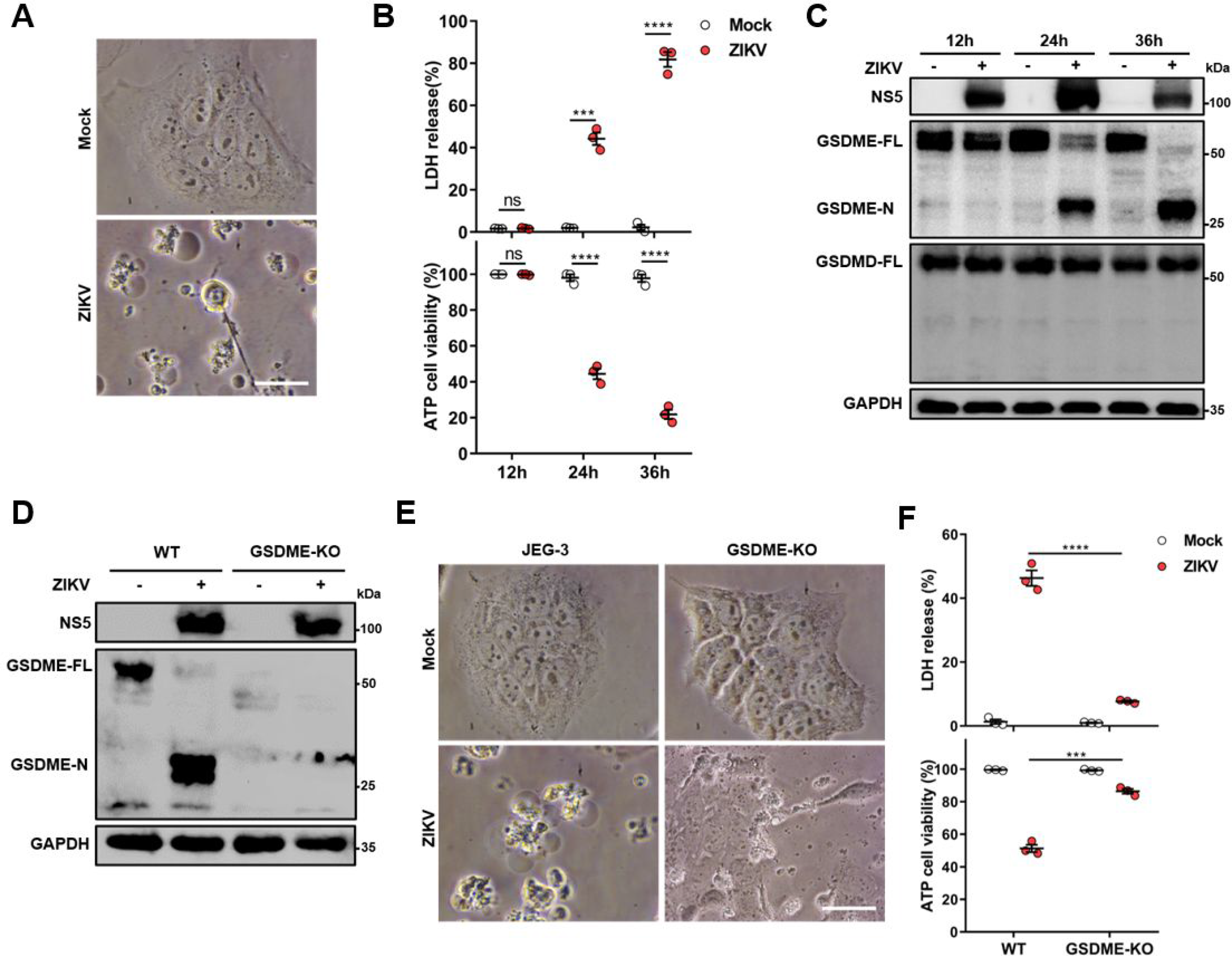
ZIKV infection induces the GSDME-mediated pyroptosis in JEG-3 cells. JEG-3 or GSDME-KO JEG-3 cells were infected with ZIKV at a MOI of 1. At indicated time post-infection, cells were subjected to microscopy, cytotoxicity and Western blot analyses. (A) JEG-3 cells were infected with ZIKV for 24 h. Representative cell morphology was shown. Scale bar, 50 μm. (B) LDH levels in supernatant and cell viability were measured at indicated time post-infection. (C) Immunoblot analyses of GSDME-FL, GSDME-N and GSDMD-FL in ZIKV-infected JEG-3 cells at indicated time post-infection. (D to F) JEG-3 and GSDME-KO JEG-3 cells were infected with ZIKV for 24 h. Immunoblot analyses of GSDME-FL and GSDME-N by Western blot (D). Representative cell morphology was shown. Scale bar, 50 μm (E). LDH levels in supernatant and cell viability were measured (F). All data are presented as the mean ± SEM of at least three independent experiments. GSDME-FL, full-length GSDME; GSDME-N, the N- terminal cleavage products of GSDME; ***, P<0,001; ****, P < 0.0001; ns, no significance.

### Both the cellular GSDME abundance and the susceptibility to ZIKV infection determine the occurrence of pyroptosis

Since there is no previous study reporting that ZIKV can induce GSDME-mediated pyroptosis, we asked whether it is a common cellular process during ZIKV infection. We infected five different human tissue-derived cell lines, including Huh-7 (liver), A549 (lung), HeLa (cervix), HEK-293T (kidney) and SH-SY5Y (neuroblasts) with ZIKV, and subsequently examined their cytopathies. It was observed that Huh-7 and A549 cells showed distinct pyroptotic cell death, elevated LDH levels, and GSDME cleavage upon ZIKV infection, whereas HeLa, HEK-293T, and SH-SY5Y cells exhibited no obvious cytopathic effects even at 72 h post-infection (Figures 2A to C). The GSDME-mediated pyroptosis is closely related to the GSDME abundance (29). We noticed that infected SH-SY5Y cells did not undergo pyroptosis despite expressing the high level of GSDME, but the relatively lower GSDME-expressing Huh-7 and A549 cells presented the different levels of pyroptosis and GSDME activation upon infection (Figures 2A-2C, and S1A), suggesting that other factors are involved in ZIKV-induced pyroptosis.

**Figure 2.**
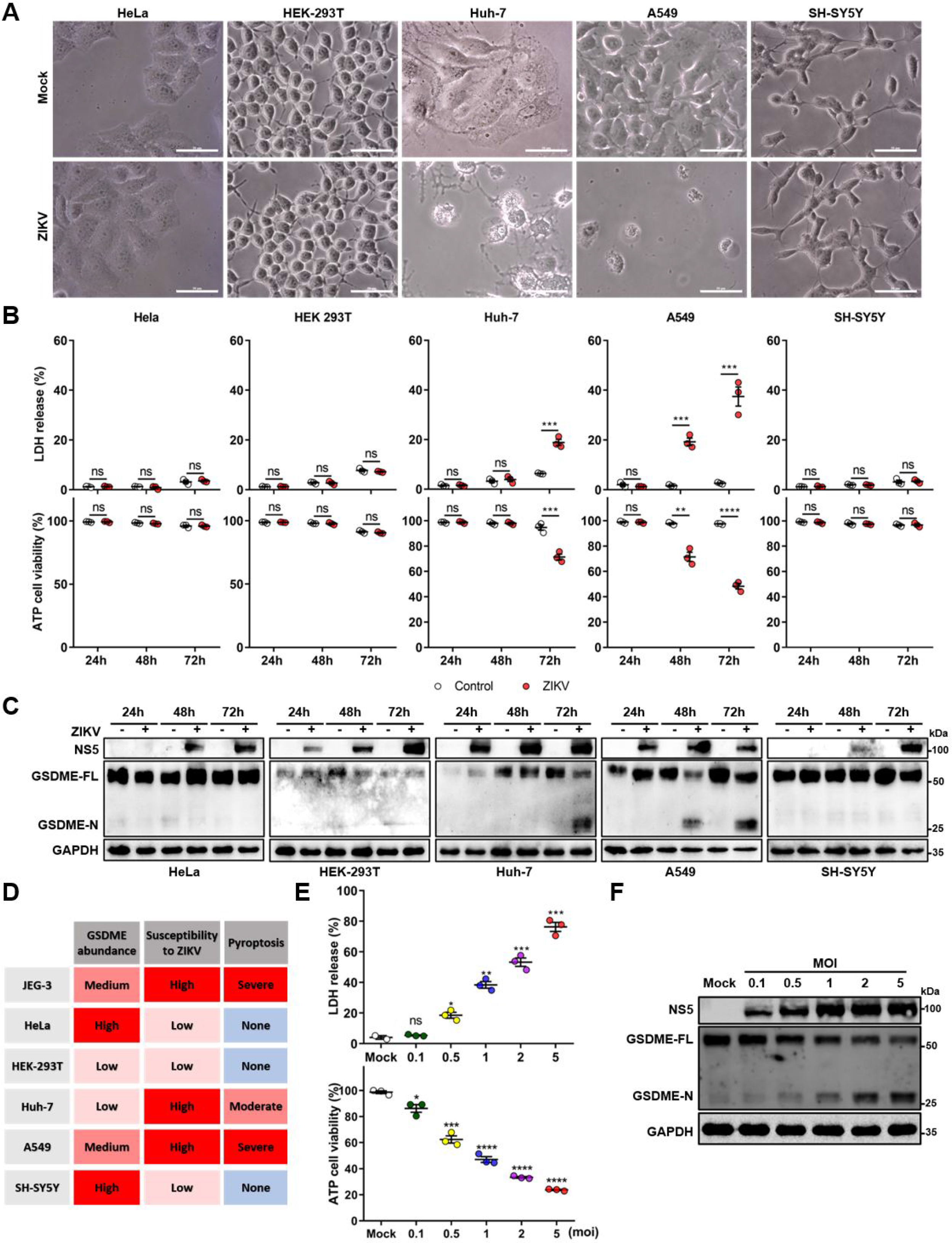
Both the cellular GSDME abundance and the susceptibility to ZIKV infection determine the occurrence of pyroptosis. (A to C) Relevant cells were infected with ZIKV at a MOI of 1. At indicated time post-infection, cells were subjected to microscopy, cytotoxicity and Western blot analyses. Phase-contrast images of ZIKV-induced pyroptotic cell death in HeLa, HEK 293T, Huh-7, A549 and SH-SY5Y cells at 72 h post-infection. Scale bar, 50 μm (A). Comparison of ATP cell viability, LDH release-based cell death (B) and GSDME cleavage (C) in ZIKV-infected relevant cells. (D) Table summarizing results shown in Figure 2A, B, C and Figure EV1 from ZIKV infection experiments with relevant cell lines. (E and F) Analyses of ATP cell viability, LDH release-based cell death (E) and GSDME cleavage (F) in JEG-3 cells at 36 hours post-infection. Unpaired t test versus mock. All data are presented as the mean ± SEM of at least three independent experiments. *, P<0.05; **, P < 0.01; ****, P<0.001; ****, P < 0.0001; ns, no significance.

In the context of infection, all these cells showed marked discrepancies in terms of susceptibility to ZIKV (36, 37). Among them, JEG-3, Huh-7 and A549 cells displayed the highest susceptibility to ZIKV infection, as assessed by the immunofluorescence assay and the plaque assay (Figure S1B and S1C). The observed positive correlation between the cellular susceptibility to infection and the increased cytopathic effects indicates that the cellular susceptibility to ZIKV infection is probably another key factor that determines the occurrence of pyroptosis (Figure 2D). We then infected JEG-3 cells with ZIKV at an increasing multiplicity of infection (MOI) to investigate whether more viral particles could lead to graver pyroptosis. At 24 h post-infection, the ZIKV-induced LDH release (Figure 2E) and GSDME activation (Figure 2F) were found to be enhanced in a virus dose-dependent manner, which implicit that the productive infection or the pathogen-associated molecular patterns of ZIKV are critical to its pathogenesis. Collectively, these results indicate that the ZIKV-GSDME-mediated pyroptosis is more likely to occur in those cells that are susceptible to ZIKV infection, and with relatively higher GSDME abundance.

### ZIKV infection activates GSDME via extrinsic apoptotic pathway

It is known that GSDME is cleaved and activated specifically by caspase-3, which consequently results in pyroptosis (28, 29). Hence, we explored the activation of caspase-3 upon ZIKV infection. At 24 h post-infection, caspase-3 underwent activation and obvious cleavage of GSDME was noticed in infected JEG-3 cells (Figure 3A). To verify if other cell death pathways are involved in the ZIKV-induced pyroptosis, ZIKV-infected JEG-3 cells were incubated with caspase-3 inhibitor (Z-DEVD-FMK), caspase-1 inhibitor (VX-765), pan-caspase inhibitor (Z-VAD-FMK) or necrosis inhibitor (GSK872). As the results are shown in Figure 3B and 3C, ZIKV infection caused the induction of pyroptotic morphological features, and the cell death was partially prevented in the presence of caspase-3 or pan-caspase inhibitors. In line with these findings, the Western blot analysis revealed that caspase-3 or pan-caspase inhibitors significantly attenuated the concomitant cleavage of caspase-3 and GSDME (Figure 3D). In contrast, the treatment of cells with caspase-1 inhibitor and necrosis inhibitor showed no effect on the cell death caused by ZIKV (Figures 3B and 3D), which suggests that necrosis and GSDMD-mediated pyroptosis were not involved in it. Altogether, these data indicate that ZIKV infection facilitates the caspase-3-dependent cleavage of GSDME to elicit pyroptotic cell death.

**Figure 3.**
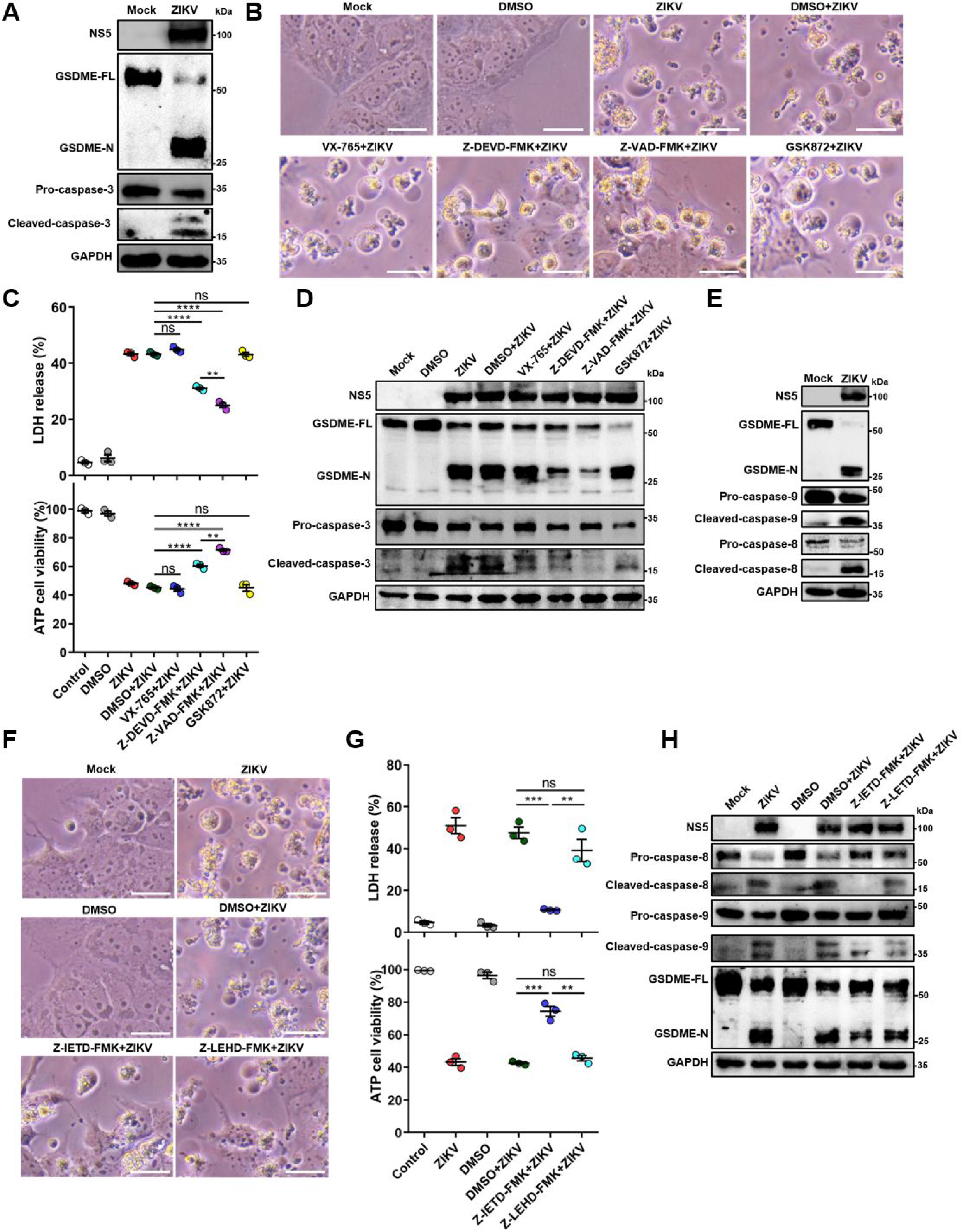
ZIKV infection activates GSDME via extrinsic apoptotic pathway. JEG-3 cells were infected with ZIKV at a MOI of 1 then subjected to indicated treatments. At 24 h post-infection, cells were subjected to microscopy, cytotoxicity and Western blot analyses. (A) Immunoblot analyses of GSDME-FL, GSDME-N, and casepase-3 in ZIKV-infected JEG-3 cells. (B to D) ZIKV-infected JEG-3 cells were incubated with either 10 μM VX-765, 25 μM Z-DEVD-FMK, 25 μM Z-VAD-FMK, or 10 μM GSK872 for 24 h and then the cells were subjected to microscopy (B), cytotoxicity (C) and Western blot analyses (D). Scale bar, 50 μm. (E) Immunoblot analyses of GSDME-FL, GSDME-N, caspase-9 and casepase-8 in ZIKV-infected JEG-3 cells. (F to H) ZIKV-infected JEG-3 cells were incubated with either 25 μM Z-IETD-FMK or 25 μM Z-LEHD-FMK for 24 h and then the cells were subjected to microscopy (F), cytotoxicity (G) and Western blot analyses (H). Scale bar, 50 μm. All data are presented as the mean ± SEM of at least three independent experiments. **, p < 0.01; ***, p < 0.001; ****, p < 0.0001; ns, no significance.

Before the caspase-3 was reported as a key regulator of the GSDME-mediated pyroptosis, it was considered an essential effector at the end of apoptotic cascades. During apoptosis, caspase-3 can be activated by intrinsic and/or extrinsic pathways: the former refers to the permeabilization of the mitochondrial membrane and the assembly of apoptosome resulting in the activation of caspase-9, and the latter involves the activation of death receptors and caspase-8 (38, 39). It has been previously demonstrated that the GSDME-dependent pyroptosis undergoes activation downstream of the intrinsic apoptotic pathway and can potentially be activated downstream of the extrinsic apoptosis pathway (17, 40, 41). However, the engagement of intrinsic and/or extrinsic pathways in order to regulate the GSDME-dependent pyroptosis during ZIKV infection, remains to be studied. To delineate the molecular mechanism by which cellular pyroptosis was instigated under ZIKV infection, we detected the activation of caspase-8 and caspase-9 in infected JEG-3 cells by the Western blot analysis. It is not unexpected to find that ZIKV infection triggered the activation of both caspase-8 and caspase-9 (Figure 3E), considering the fact that various biological processes are triggered during ZIKV infection. Next, the ZIKV-infected cells incubated with caspase-8 inhibitor (Z-IETD-FMK) or caspase-9 inhibitor (Z-LEHD-FMK), and subsequently were subjected to the analysis of pyroptotic cell parameters. It was found that caspase-8 inhibitor remarkably inhibited the cell membrane rupture, LDH release, and GSDME cleavage in infected cells (Figures 3F and3 H). Interestingly, the caspase-8 inhibitor treatment also reduced the activation of caspase-9, and the caspase-9 inhibitor treatment slightly inhibited the activation of GSDME without exerting any impact on the ZIKV-induced pyroptotic cell death (Figures 3F and 3H). These findings urged us to reflect on whether caspase-8 was activated before caspase-9 in the GSDME-mediated signaling pathway. As recent studies demonstrate that the GSDME-N can permeabilize the mitochondrial membrane and activate the intrinsic apoptotic pathway (42), it implies the possibility that during the activation of GSDME upon ZIKV infection the caspase-8-associated extrinsic apoptosis may have activated first, which consequently targets mitochondria to activate the caspase-9-associated intrinsic apoptosis, thereby creating a self-amplifying feed-forward loop through consecutively activating caspase-3. Given that the activation of death receptors upon ligand binding is a crucial step in the extrinsic apoptosis pathway, we then evaluated the influence of ZIKV infection on the expression of tumor necrosis factor α (TNF-α). ZIKV infection significantly enhanced the expression of TNF-α in infected JEG-3 cells, as determined by quantitative real-time reverse transcription PCR (qRT-PCR) (Figure S2). Despite elusive mechanistic underpinnings, these data suggest that the extrinsic apoptotic pathway plays a key role in mediating the ZIKV-induced pyroptosis, and that the potential crosstalk between pyroptosis and apoptosis during the ZIKV pathogenesis could not be neglected.

### The genomic RNA of ZIKV activates the GSDME-dependent pyroptosis through RIG-I pathway

In order to explore which part of the ZIKV is responsible for inducing pyroptosis, we first infected JEG-3 cells with ZIKV or UV-inactivated ZIKV to test the contribution of viral structural proteins in inducing the pyroptosis, and found that ZIKV structural proteins are not functional pyroptosis agonists (Figures S3A and S3B). Next, we transfected JEG-3 cells with individual viral non-structural (NS) proteins fused with the N-terminal 3XFlag-tag to find which NS proteins are capable of activating pyroptosis. The results showed that all NS proteins had no effect on pyroptosis (Figures 4A and S3C), hence guiding us to speculate that the viral RNA may participate in orchestrating this phenomenon.

**Figure 4.**
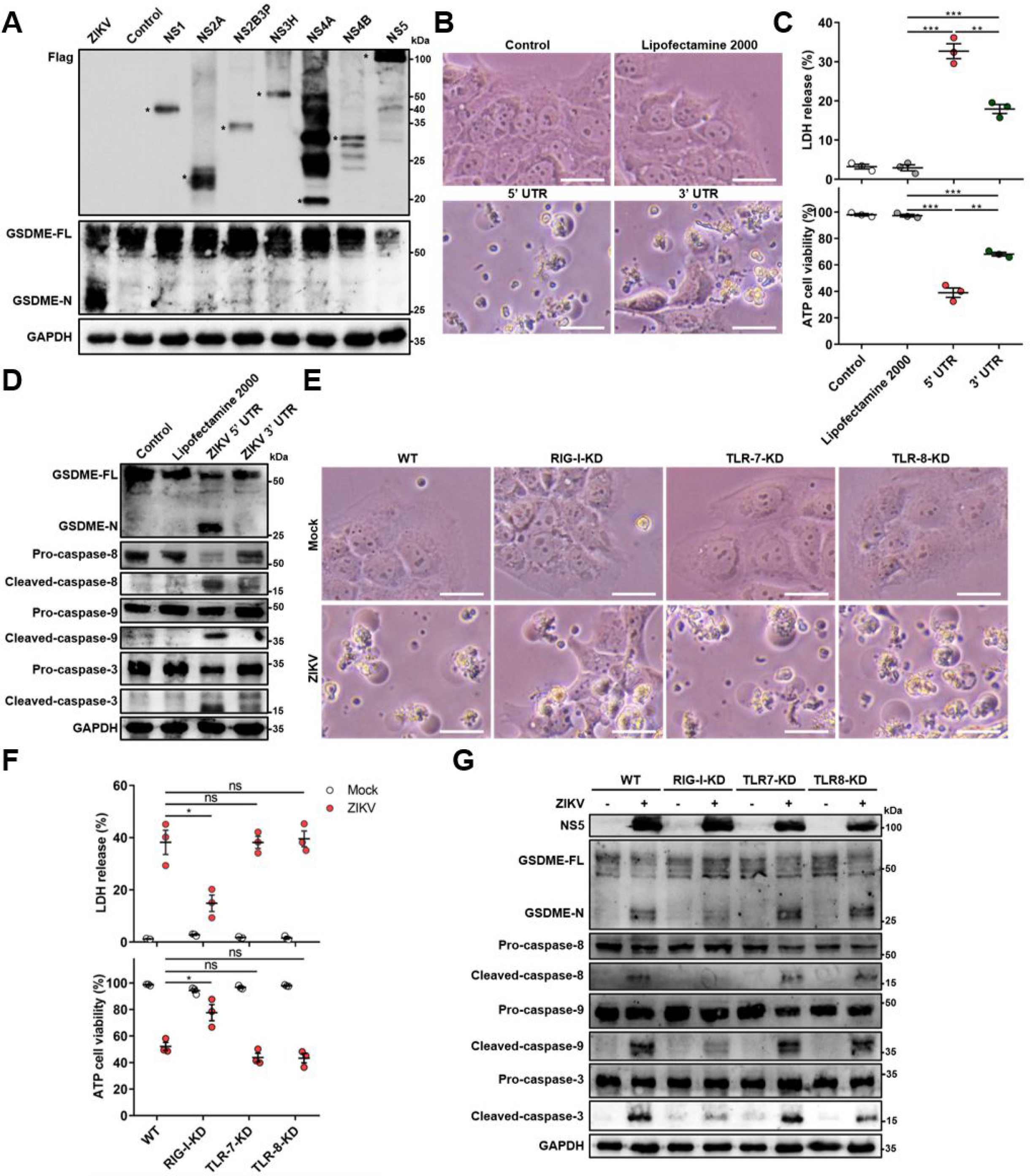
The genomic RNA of ZIKV activates the GSDME-dependent pyroptosis through RIG-I-caspase-8-caspase-3 pathway. (A) JEG-3 cells were seeded in 6-well plate, and transfected with 2 μg indicated plasmids. At 24 h post-transfection, cells were subjected to analyze for GSDME-FL, GSDME-N and Flag by Western blot. Asterisks indicate specific bands. (B to D) JEG-3 cells were seeded in 6-well plate and were transfected with 1 μg ZIKV 5’ UTR or 3’ UTR. At 24 h post-transfection, cells were subjected to microscopy (B), cytotoxicity (C) and Western blot analyses. Scale bar, 50 μm. (E to G) Relevant cell lines were infected with ZIKV at a MOI of 1. At 24 h post-infection, cells were subjected to microscopy (E), cytotoxicity (F) and analyzed for GSDME-FL, GSDME-N, caspase-8, caspase-9 and caspase-3 by Western blot (G). Scale bar, 50 μm. All data are presented as the mean ± SEM of at least three independent experiments. **, p < 0.01; ***, p < 0.001; ****, p < 0.0001; ns, no significance.

For this purpose, we obtained the 5’ and 3’ untranslated region (UTR) RNA of the ZIKV through the in vitro transcription assay to mimic the viral genome, and subsequently transfected them into JEG-3 cells. As the results shown in Figures 4B and 4C, the UTRs induced the distinct pyroptotic cell death and LDH release. In consistent with these results, the Western blot analysis exhibited that the ZIKV UTRs caused the activation of GSDME via the same signaling pathway as induced by ZIKV infection (Figure 4D). It is worth noting that, compared with the ZIKV 5’UTR, the 3’ UTR caused relatively a weaker effect on inducing pyroptosis (Figures 4B and 4C), hinting that the recognition of specific viral RNA motifs may aid the GSDME-dependent pyroptosis.

Similar to other flaviviruses, ZIKV contains a single-stranded (ss), positive-sense RNA genome. Upon infection of the host cell by RNA viruses, cellular pattern recognition receptors (PRRs) play a decisive role in detecting viral RNAs and restricting viruses’ replication. Among the PRRs, the retinoic acid-inducible gene I (RIG-I) and Toll-like receptor (TLR) 7 and 8 are responsible for sensing the viral ssRNA genome (43, 44). Based on the above-mentioned results that the ZIKV RNA genome stimulates pyroptosis, whether RIG-I, TLR7 or TLR8 contributes to the onset of this process was evaluated. The RIG-I-knockdown (KD), TLR7-KD, or TLR8-KD JEG-3 cell lines were generated using the CRISPR/Cas9 system (Figure S4A). ZIKV infection of engineered or wild-type (WT) cells revealed that the KD of RIG-I, but not TLR7 and TLR8, conspicuously attenuated the ZIKV-induced pyroptotic cell death and GSDME cleavage, when compared to that in the WT cells (Figures 4E and 4F). In addition, the KD of RIG-I suppressed the ZIKV-induced production of TNF-α, and the activation of pyroptosis caused by ZIKV 5’ UTR (Figures S4B and S4C), suggesting that RIG-I-mediated recognition of ZIKV genome is important for launching pyroptosis. Taken together, these results manifest that the ZIKV genome acts as a pivotal factor in inducing pyroptosis through RIG-I pathway.

### ZIKV infection of pregnant immunocompetent WT C57BL/6N mice results in placental pyroptosis and CZS

To verify whether ZIKV infection lead to pyroptosis of placental tissue in vivo, WT C57BL/6N mice were infected intravenously with 1 × 10^6^ PFU of ZIKV French Polynesia strain H/PF/2013 or with an equal volume of Vero cell culture supernatant at embryonic day 9.5 (E9.5) of pregnancy. At E16.5, mice were intravenously administrated with propidium iodide (PI) to evaluate cell viability, and were sacrificed 15 min later. Placentas and individual fetuses were evaluated for morphological appearance (scheme outlined in Figure 5A). Notably, most of dams infected with ZIKV showed abnormal pregnancies (Figure S5A), and the fetuses of mock-infected mice displayed apparent differences compared to the ZIKV-infected mice, which contained fetuses showing a variable degree of CZS (Figure 5B). Among all 14 ZIKV-infected dams, only 2 dams displayed normal pregnancies. The remaining 12 dams exhibited abnormal pregnancies, with 4 dams containing only placental residues and embryonic debris as all fetuses had undergone resorption. Other 8 infected dams carried at least one or more placental residues and morphologically abnormal fetuses. Out of 109 implantation sites carried by all 14 infected dams, 67 (61%) sites were affected, among which 52 showed fetal resorptions and 15 showed growth restriction and malformation (Figures S5C and S5B). These data indicate that ZIKV infection indeed leads to CZS in our experimental model.

**Figure 5.**
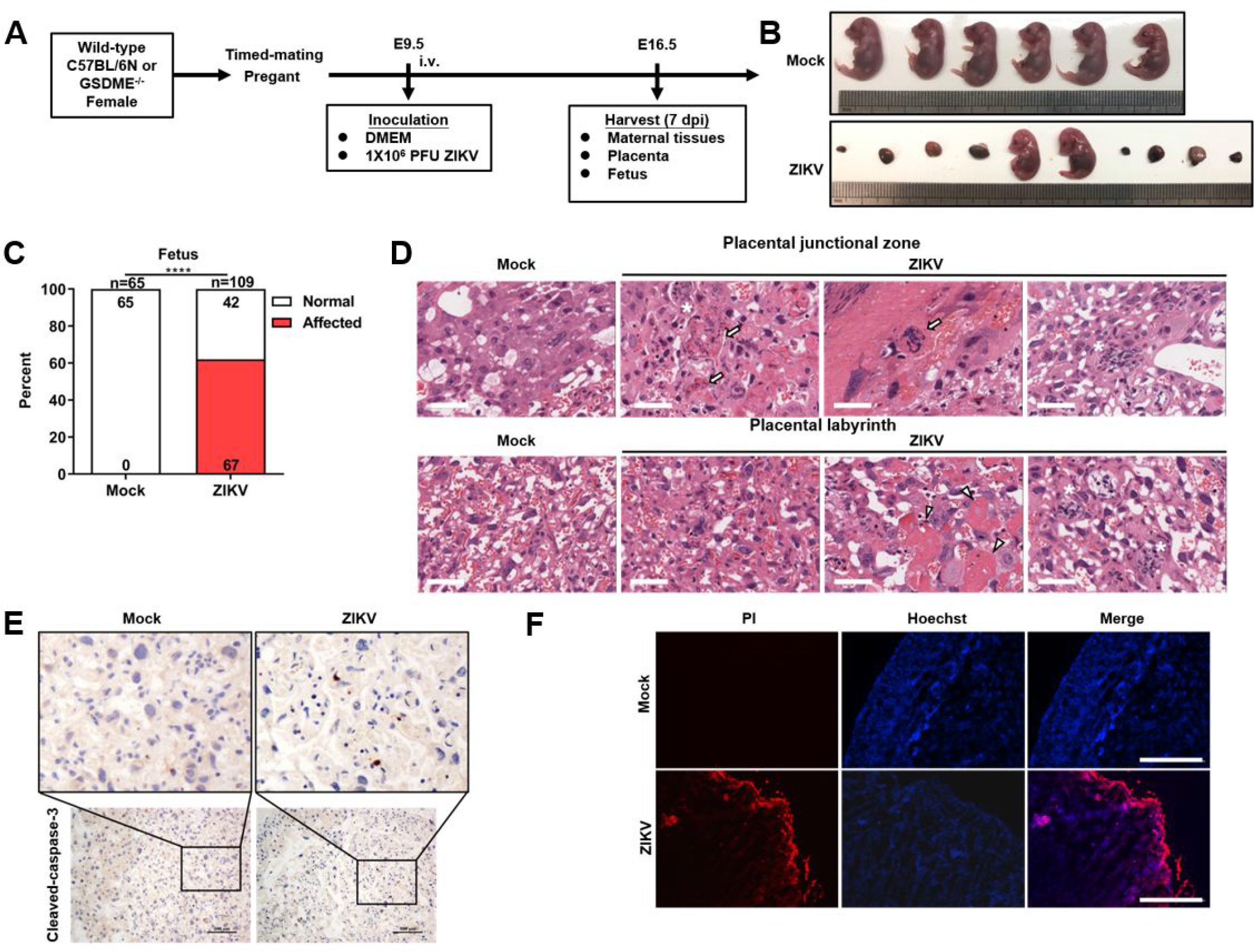
ZIKV infection of pregnant immunocompetent WT C57BL/6N mice results in placental pyroptosis and CZS. Pregnant dams at E9.5 were infected with 1 × 106 PFU of ZIKV by intravenous route. Mock mice were inoculated with Vero cell culture supernatant. Mice were sacrificed 7 days post-infection at E16.5, and maternal tissues, placentas and fetuses were harvested and analyzed. (A) ) Scheme of infection and the follow up analyses. (B) Representative images of fetuses at E16.5 are shown. (C) Impact of ZIKV infection on fetuses at E16.5. The percentage of fetuses that were affected (i.e. had undergone resorption, or exhibited any sign of growth restriction, malformation) is shown. Numbers on bars indicate normal fetuses (top) or affected fetuses (bottom). (D) Representative hematoxylin and eosin staining showed pathological features of placentas at E16.5. The asterisks indicate abnormal spheroid structure. Arrows indicate necrotic trophoblast cells. Arrowheads indicate thrombi. Scale bar, 50 μm. (E) Representative images of immunohistochemistry of placenta sections stained for Cleaved-caspase-3. Scale bar, 100 μm. (F) Propidium iodide was intravenously injected into the mice before assay. Representative placenta section images are shown. Scale bar, 400 μm. Number of samples in each group is listed. Data for all panels are pooled from 3–5 independent experiments. For (C), significance was determined by Fisher’s exact test. Data shown are median with interquartile range. ****, p < 0.0001; ns, no significance.

Furthermore, the general histological examination of the placentas was conducted in order to observe the pathological changes associated with ZIKV infection. Mock-infected placentas presented normal features of the embryo-derived junctional and labyrinth zone, a place where the nutrient exchange occurs (Figure 5D). In infected placentas, obvious abnormal morphological changes were noticed, including hyperplasia, necrosis and thrombi. Specifically, the hyperplastic trophoblast labyrinth showed denser cellularity and less vascular spaces, suggesting that the placental and embryonic blood circulation may have been compromised. Otherwise, abnormal spheroid structures were found in the junctional and labyrinth zone, which contained necrotic trophoblast cells, and thrombi were observed in the labyrinth zone. Hence, these observed changes indicate that ZIKV infection contributes to placenta pathology.

As previous studies reported that the placental pathology during ZIKV infection corresponds to adverse fetal outcomes (5, 15), we asked whether the GSDME-mediated placental pyroptosis is involved in regulating this phenomenon. We first tested the activation of caspase-3 (a key signaling molecule upstream of GSDME), and the immunohistochemistry assay showed strong activation of it in the trophoblast labyrinth (Figure 5E). Moreover, we conducted the PI staining assay by considering the fact that pyroptotic cells are permeable to small molecular weight, membrane-impermeable dyes such as 7-aminoactinomycin D, ethidium bromide, and PI (20). No PI-positive cells were visualized in the placentas of mock-infected dams. In contrast, in the placentas of ZIKV-infected dams, we detected PI-positive signal in the decidua and labyrinth zone, suggesting the occurrence of pyroptosis (Figure 5F). Taken together, these findings emphasize the importance of ZIKV-induced placental injury in developing CZS, and establish a link between placental injury and pyroptosis.

### GSDME contributes to the ZIKV-induced CZS through mediating placental pyroptosis

To determine the role of GSDME in contributing to the ZIKV-induced placental damage and adverse fetal outcomes, Gsdme^-/-^ mice were used. WT or Gsdme^-/-^ mice were infected intravenously with 1 × 10^6^ PFU of ZIKV French Polynesia strain H/PF/2013 or with an equal volume of Vero cell culture supernatant at E9.5 of pregnancy, and were sacrificed 7 days later at E16.5 (scheme outlined in Figure 5A). Although ZIKV-infected Gsdme^-/-^ dams also showed a certain degree of abnormal pregnancies, the number and severity of affected dams and fetuses were markedly decreased compared to the WT dams (Figures 6A-6B, and S6). Among all 10 ZIKV-infected Gsdme^-/-^ dams, 4 dams showed abnormal pregnancies with a mixture of placental residues and stunted embryonic growth. The case in which all fetuses were resorbed in a dam vanished, and all fetuses carried by the rest of 6 dams showed no morphological abnormalities. While 67 (61%) fetuses carried by the ZIKV-infected WT dams were affected, the affected fetuses in the ZIKV-infected Gsdme^-/-^ dams were only 18 (20%), among which 14 showed fetal resorptions and 4 showed growth restriction and malformation. Thus, these data indicate that GSDME is associated with ZIKV-induced adverse fetal outcomes.

**Figure 6.**
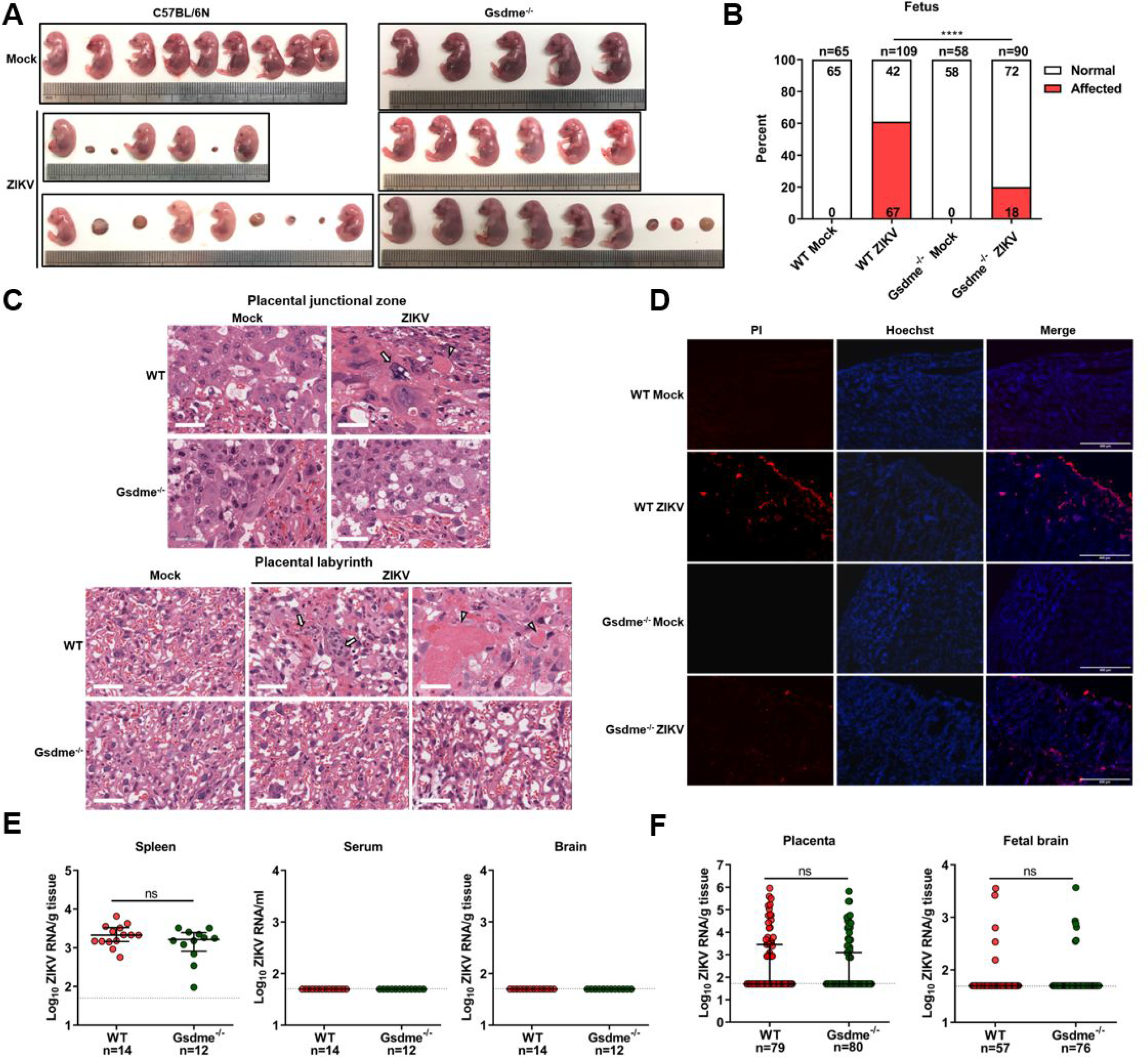
GSDME contributes to the ZIKV-induced CZS through mediating placental pyroptosis. Scheme of infection and analyses were shown in Figure 5A. The WT Mock and WT ZIKV groups were the same groups as shown in Figure 5. (A) ) Representative images of fetuses from WT and Gsdme^-/-^ dams at E16.5 are shown. (B) Impact of ZIKV infection on fetuses from WT and Gsdme^-/-^ dams at E16.5. The percentage of fetuses that were affected (i.e. had undergone resorption, or exhibited any sign of growth restriction, malformation) is shown. Numbers on bars indicate normal fetuses (top) or affected fetuses (bottom). (C) Representative hematoxylin and eosin staining showed pathological features of placentas at E16.5. The asterisks indicate abnormal spheroid structure. Arrows indicate necrotic trophoblast cells. Arrowheads indicate thrombi. Scale bar, 50 μm. (D) Propidium iodide was intravenously injected into the mice before assay. Representative placenta section images are shown. Scale bar, 400 μm. (E) ZIKV RNA levels of maternal spleens, serum and brains of WT and Gsdme^-/-^ dams infected with ZIKV. (F) ZIKV RNA levels of all placentas and fetal heads carried by WT and Gsdme^-/-^ dams infected with ZIKV. Number of samples in each group is listed. Data for all panels are pooled from 3–5 independent experiments. For (B), significance was determined by Fisher’s exact test. For (E) and (F), the Mann-Whitney test was used to calculate significance. Data shown are median with interquartile range, and the dotted line depicts the limit of detection. ****, p < 0.0001; ns, no significance.

We next performed the histological examination of placentas regardless of whether their corresponding fetuses were affected or not. Compared to the placentas from ZIKV-infected WT dams, trophoblastic hyperplasia, reduced vascular spaces, abnormal spheroid structures, and necrotic cells were not seen in the placentas from ZIKV-infected Gsdme^-/-^ dams (Figure 6C). In addition, the PI staining assay revealed relatively fewer PI-positive signals in the placentas from ZIKV-infected Gsdme^-/-^ dams, implying that the deficiency of GSDME inhibits the pyroptosis of placentas (Figure 6D).

In order to rule out the possibility that the variable infection status of ZIKV in WT and Gsdme^-/-^ mice may confer variable fetal outcomes, we measured the ZIKV RNA levels in maternal tissues (spleen, brain, and serum), placentas, and fetuses (fetal brain). Comparable ZIKV RNA levels were detected in the spleens of these two types of mice, but the viral RNA remained undetectable in their brains and serum (Figure 6E). No significant differences were found in viral copies and infection ratio between the placentas and fetal brains from ZIKV-infected WT and Gsdme^-/-^ dams (Figure 6F). Thereby, based on these data, we conclude that maternal GSDME does not affect ZIKV infection, but the GSDME in placenta promotes CZS by mediating placental pyroptosis.

## Discussion

In this study, we identified a novel mechanism that ZIKV utilizes GSDME to induce the pyroptosis of placental cells in vitro and in vivo, which consequently associates with adverse fetal outcomes in infected mice. These data established a potential relevance of the placental pyroptotic process with the ZIKV-associated adverse fetal outcomes, thus suggesting that placental pyroptosis is a vital feature of ZIKV pathogenesis.

Previously, it has been reported that ZIKV infection activates inflammatory pathways and facilitates the secretion of pro-inflammatory cytokine interleukin-1β (IL-1β) in vitro and in vivo (45, 46). ZIKV NS5 and NS1 proteins, by interacting with NLRP3 and by stabilizing caspase-1, respectively, promote the NLRP3 inflammasome assembly/activation to augment the IL-1β secretion (47, 48). Since the NLRP3 inflammasome functions upstream of pyroptosis executor GSDMD and IL-1β is passively released during cell lysis (20), it can be assumed that pyroptosis may confer ZIKV pathogenesis. Interestingly, other than the mechanisms mentioned above, our results demonstrate that ZIKV triggered pyroptosis through activating GSDME in placenta. So far, researches on GSDME have mainly focused on its anti-cancer effects, but little is known about its function in the pathogenesis of virus. Our research not only identifies for the first time that GSDME can be activated by ZIKV, but also hints the important role of GSDME in viral pathogenesis. Different from a recent study indicating that ZIKV induces the GSDMD-mediated pyroptosis of neural progenitor cells (33), our data revealed a previously unrecognized mechanism of ZIKV-induced fetal adverse outcomes, which is indirectly caused by GSDME-mediated pyroptosis. Unlike the GSDMD-mediated pyroptosis driven by caspase-1 and caspase-4/5/11, we found that GSDME undergoes activation by caspase-8 and caspase-3, which is partially consistent with published studies, wherein GSDME was subjected to be cleaved and activated by caspase-3 and granzyme B (22, 23, 29, 41).

In our study, pyroptosis was not found as a universal phenomenon upon ZIKV infection, despite the fact that ZIKV can infect multiple cell lines. For instance, ZIKV-infected HeLa, HEK293T, and SH-SY5Y cells exhibited no apparent pyroptotic morphological changes, albeit some of them express GSDME in abundance. By evaluating the susceptibility of low GSDME-expressing HEK-293T and Huh-7 cells to ZIKV infection, we found that the infection susceptibility is an additional decisive factor, as Huh-7 cells, which are highly susceptible to ZIKV, were prone to pyroptosis (Figures 2A-2C and EV1). These findings not only highlight the important role of GSDME in pathogenesis of ZIKV, but also raise the question of whether other ZIKV-induced symptoms are associated with GSDME-mediated pyroptosis, which could be alluring to investigate in future studies.

ZIKV-induced apoptosis has been widely described in the central nervous system and the female reproductive system, in which caspase-3 plays a crucial role (5, 17). Given that caspase-3 is downstream of caspase-8 and caspase-9, which individually correspond to extrinsic and intrinsic apoptosis pathways, caspase-8- and caspase-9-specific inhibitors were used to assess their respective effects on activating the caspase-3-GSDME-mediated pyroptosis during ZIKV infection of JEG-3 cells. Inhibition of caspase-8 attenuated the pyroptotic cell features and GSDME cleavage in infected cells, whereas caspase-9 inhibition caused no such effects in a significant manner. Our observation of increased TNF-α production in ZIKV-infected cells also supports this conclusion, as TNF-α is a vital ligand to commence the caspase-8-mediated signaling (49). Caspase-8 is a multifunctional effector protein related to various signaling pathways, and acts as a molecular switch for apoptosis, necrosis, and pyroptosis (50). In the case of pyroptosis, caspase-8 is capable of inducing the GSDMD- and GSDME-mediated pyroptosis in murine macrophages (51-53). However, our data showed that GSDMD was not activated by caspase-8 in ZIKV-infected JEG-3 cells, inferring the possibility that some specific conditions are required for inducing the GSDMD-mediated pyroptosis. In addition, caspase-8 inhibits necrosis regulated by receptor-interacting serine/threonine kinase 3 (49), which fortifies our conclusion that necrosis did not occur in ZIKV-infected JEG-3 cells.

During the viral infection of host cells, PRRs are vital for detecting the viral genomes and limiting the spread of viruses (54). For instance, RIG-I and TLR7 recognize the genomic RNAs of hepatitis C and West Nile viruses, respectively, to induce host defense m(55, 56). Based on our findings that the ZIKV RNA genome, rather than the ZIKV-encoded proteins, spurs pyroptosis, we asked if PPRs, including TLR7, TLR8, and RIG-I were involved in the occurrence of this cell death process. Using the CRISPR-Cas9-engineered respective KD JEG-3 cells, we observed the necessity of RIG-I to induce the GSDME-mediated pyroptosis via recognizing the ZIKV RNA genome, with a piece of further evidence that the ZIKV 5’UTR was more effective than the 3’UTR in initiating pyroptosis. These data are in concordant with a recent study which demonstrated that RIG-I recognizes the ZIKV 5’UTR in human cells and the enrichment of viral RNA on RIG-I decreases along with the genome toward the 3’UTR (57), hence proposing a possibility that the activation degree of RIG-I determines the outcome of pyroptosis.

Several studies by using immunocompromised mouse models have demonstrated the detection of ZIKV in the placentas and embryos, resulted in trophoblast apoptosis, vascular damage, loss of placental structure, and reduced neonatal brain cortical thickness (5, 15). Under natural conditions, mice are resistant to ZIKV due to the activation of type-I interferon and research has illustrated that type-I interferon signaling may play a vital role in mediating fetal demise after ZIKV infection by causing abnormal placental development (18), implying that maternal immune is response to undertake the antiviral role and also promotes the occurrence of pathologies. In this context, to simulate the pathological changes triggered by ZIKV-infection in the natural state as much as possible, we employed immunocompetent mice for experiments. It is worth noting that regardless of whether placental lesions and fetal resorption occurred or not, the ZIKV RNA was infrequently detected, which suggests that placental function may be affected under a negligible quantity of viral particles, or the virus had been eliminated at this stage. As the placenta acts as a crucial tissue between the gravida and embryo, preventing pathogen transmission during pregnancy and mediating the exchange of nutrients imply that any inappreciable placental damage can lead to devastating consequences.

In summary, our study identified a novel mechanism of ZIKV pathogenesis that causes placental pyroptosis in vitro and in vivo, triggered upon the recognition of the viral genome by RIG-I followed by the caspase-8- and caspase-3-mediated activation of pyroptosis executor GSDME in placental cells. The ablation of GSDME in infected pregnant mice attenuated placental pyroptosis and adverse fetal outcomes. These findings may provide opportunities to establish therapies that might halt placental damage and subsequent adverse fetal outcomes during ZIKV infection.

## Materials and Methods

### Ethics statement

All animal studies were conducted in strict accordance with the Guide for Care and Use of Laboratory Animals of Laboratory Animal Centre, Huazhong Agriculture University, and all experiments conform to the relevant regulatory standards. The experiments and protocols were approved by the Animal Management and Ethics Committee of Huazhong Agriculture University (Assurance number HZAUMO-2020-0053).

### Reagents and antibodies

Mouse monoclonal anti-ZIKV NS5 antibody was generated in our lab. Rabbit monoclonal anti-GSDME (ab215191) antibody was purchased from Abcam. Mouse monoclonal anti-GAPDH (AC002), mouse monoclonal anti-Flag (AE005), rabbit monoclonal anti-TLR7 (A19126), rabbit polyclonal anti-TLR8 (A12906), and rabbit monoclonal anti-caspase-3 (A19654) antibodies were purchased from ABclonal Technology. Rabbit polyclonal anti-GSDMD (96458>), mouse monoclonal anti-caspase-8 (9746), and rabbit polyclonal anti-caspase-9 (9502) antibodies were purchased from Cell Signaling Technology. Rabbit polyclonal anti-RIG-I (20566-1-AP) was purchased from Proteintech. Horseradish peroxidase-labeled goat anti-mouse (AS003) or anti-rabbit (AS014) secondary antibodies were obtained from ABclonal Technology.

The chemical reagents DAPI (D8417), PI (P4170) and Hoechst (B2261) were purchased from Sigma-Aldrich. The inhibitors specific to pan-caspase (zVAD-FMK), caspase-1 (VX-765), caspase-3 (Z-DEVD-FMK), caspase-8 (Z-IETD-FMK), caspase-9 (Z-LETD-FMK), and necrosis (GSK-872) were purchased from Selleck.

### Cell culture, viruses and plaque assay

All cell lines were obtained from the American Type Culture Collection (ATCC). Human embryonic kidney (HEK-293T), human alveolar epithelial adenocarcinoma (A549), human hepatocellular carcinoma (Huh-7), and African green monkey kidney (Vero) cells were cultured in Dulbecco’s modifed Eagle’s medium (DMEM; Sigma) supplemented with 100 U/ml penicillin, 100 g/ml streptomycin, and 10% fetal bovine serum (FBS, GIBCO). Aedes albopictus mosquito (C6/36) and human cervical cancer (HeLa) cells were cultured in RPMI-1640 medium (Hyclone) supplemented with 10% FBS and 2 mM L-glutamine. Human neuroblastoma (SH-SY5Y) and human choriocarcinoma/trophoblastic cancer (JEG-3) cells were grown in DMEM supplemented with 10% FBS and MEM Non-essential Amino Acid Solution (Sigma). C6/36 cells were grown at 28°C in a 5% CO2 incubator, whereas all other cells were grown at 37°C in a 5% CO2 incubator. The ZIKV strain H/PF/2013 was propagated in Vero cells and were titrated by plaque assays on Vero cells. In brief, relevant cells were inoculated with ZIKV at a MOI of 0.1, and the culture supernatants were harvested at 12-72 h post-infection. The supernatants were then serially diluted and inoculated onto the monolayers of Vero cells. After 1 h of absorption, cells were washed with serum-free DMEM and cultured in DMEM containing 3% FBS and 1.5% sodium carboxymethyl cellulose (Sigma-Aldrich). Visible plaques were counted, and viral titers were calculated after 5 days of incubation.

### Cytotoxicity assay

Cell death was assessed by the LDH assay using the CytoTox 96 Non-Radioactive Cytotoxicity Assay kit (Promega). Cell viability was determined by the CellTiter-Glo Luminescent Cell Viability Assay kit (Promega). All the relevant experiments were performed following the manufacturer’s instructions.

### Western blot analysis

Total cell lysates were prepared using the RIPA buffer (Sigma) containing protease inhibitors (Roche). After sonication, the protein concentration in each sample was determined by using the BCA protein assay kit (Thermo Fisher Scientific) and boiled at 95°C for 10 min. Equivalent amounts of protein samples were separated by SDS-PAGE and electroblotted onto a polyvinylidene fluoride membrane (Roche) using a Mini Trans-Blot Cell (Bio-Rad). The membranes were blocked at room temperature (RT) for 2 h in PBS containing 3% bovine serum albumin (BSA), followed by incubation with the indicated primary antibodies overnight at 4°C. After washing three times with TBS-Tween (50 mM Tris-HCl, 150 mM NaCl, and 0.1% [v/v] Tween 20, pH 7.4), the membranes were incubated with secondary antibodies at RT for 45 min. Finally, the membranes were visualized with a chemiluminescence system (Bio-Rad) after three times of wash.

### Microscopy and immunofluorescence analysis

To examine the morphology of pyroptotic cells, relevant cells were seeded in 6-well plates at ∼40% confluency and then subjected to indicated treatments. Phase contrast cell images were captured by the NIKON Ti-U microscope.

For immunofluorescence analysis, relevant cells were seeded in 12-well plates and infected with ZIKV at a MOI of 0.1. At indicated time post-infection, cells were fixed with pre-cold methanol for 10 min at −20°C. Subsequently, cells were washed three times by PBS and were blocked in PBS containing 1% BSA for 1 h at RT. Thereafter, cells were incubated with anti-ZIKV NS5 antibody for 2 h at RT., followed by washing three times with PBS and incubation with Alexa Fluor 488-conjugated goat anti-mouse (Invitrogen) for 45 min. Cells were then washed and incubated with DAPI for another 10 min at RT. After final washing, cells were visualized using the EVOS FL auto (Thermo Fisher Scientific).

### Plasmid construction and transfection

Using the ZIKV cDNA as template, NS1, NS2A, NS2B3P, NS3H, NS4A, NS4B, and NS5 were separately cloned into the p3XFLAG-CMV-10 using the PCR/restriction digest-based cloning method, and were finally verified by sequencing. Transfections were performed using the Lipofectamine 2000 (Invitrogen) according to the manufacturer’s instructions.

### CRISPR-Cas9 knockdown and knockout cells

Two guide RNAs (gRNAs) target to each desired gene were cloned into the lentiviral vector lentiCRISPR v2. About 800 ng lentiviral vector, 400 ng packaging plasmid pMD2.G, and 800 ng pSPAX2 were co-transfected into HEK 293T cells using the FuGENE HD Transfection Reagent (Promega) according to the manufacturer’s instructions. At 48 h post-transfection, viral supernatants were collected, and then inoculated to 4 × 105 JEG-3 cells for another 48 h. For RIG-I, TLR-7, and TLR-8 knockdown in JEG-3 cells, the gRNA-expressing cells were selected with 1.5 μg/ml puromycin, and then plated into 12-well plates for further experiments. For GSDME knockout in JEG-3 cells, the gRNA-expressing cells were selected with 1.5 μg/ml puromycin following the cell sorting using a flow cytometry system. The single-cell clones were cultured in 96-well plates for another 10 days or longer. The immunoblotting was used to screen for GSDME-deficient clones and to verify the knockdown efficiency. The genotyping of the knockout cells was determined by sequencing.

The sequences of gRNAs were as follows: gRNA-GSDME-1, 5’-TAAGTTACAGCTTCTAAGTC-3’; gRNA-GSDME-2, 5’-CAGTTTTTATCCCTCACCCT-3’; gRNA-RIG-I-1, 5’-GGATTATATCCGGAAGACCC-3’; gRNA-RIG-I-2, 5’-TCCTGAGCTACATGGCCCCC-3’; gRNA-TLR-7-1, 5’-GGTGAGGTTCGTGGTGTTCG-3’; gRNA-TLR-7-2, 5’-CCTGCGGTATCTCTAGTAGC-3’; gRNA-TLR-8-1, 5’-GTGCAGCAATCGTCGACTAC-3’; gRNA-TLR-8-2, 5’-AATCCCGGTATACAATCAAA-3’.

### Mice and infections

C57BL/6N WT mice were purchased from Vital River Laboratory Animal Technology Co. Gsdme^−/−^ mice were kindly provided by Prof. Feng Shao (National Institute of Biological Sciences, Beijing). Mice were set up for timed matings, and were injected intravenously with 50 μL Vero cell culture supernatant or 1 × 10^6^ PFU of ZIKV in a volume of 50 μL at the E9.5. Mice were then sacrificed at E16.5, and placentas, fetuses, and maternal tissues were harvested for the subsequent analyses.

### Measurement of viral burden

ZIKV-infected pregnant mice were euthanized at E16.5. Maternal tissues, fetuses, and their corresponding placentas were harvested. Samples were weighed and homogenized with stainless steel beads in 1mL of DEME supplemented with 2% heat-inactivated FBS. All homogenized tissues were stored at -80℃ until virus titration was performed. Some placentas were divided equally into two parts, and one part was fixed in 4% paraformaldehyde for histological examination.

Total RNA in homogenized samples and serum were extracted using the TRIzol Reagent (Invitrogen) according to the manufacturer’s instructions, and were reverse-transcribed to cDNA with random hexamers. ZIKV RNA levels were determined by quantitative real-time PCR on a 7500 Real-time PCR system (Applied Biosystems). Viral burden was expressed on a log10 scale as viral RNA equivalents per g or per mL after comparison with a standard curve produced using the serial 10-fold dilutions of ZIKV RNA. The ZIKV-specific primer set and probe were used as published previously (58) : forward primer, 5’-CCGCTGCCCAACACAAG-3’; reverse primer, 5’-CCACTAACGTTCTTTTGCAGACAT-3’; and Probe, 5’-/56-FAM/AGCCTACCT/ZEN/TGACAAGCAATCAGACACTCAA/3IABkFQ/-3’ (Sangon).

### Histological and immunohistochemistry staining

For histological staining, placentas were fixed in 4% paraformaldehyde overnight at 4°C and embedded in paraffin. At least three placentas from different litters were sectioned and stained with hematoxylin and eosin.

For immunohistochemistry staining, ∼5 μm thick paraffin sections were placed in 3% H2O2 for 30 min to quench the endogenous peroxidase activity, and then the sectioned slides were incubated in citrate buffer at 96°C for 30 min for antigen retrieval. After washing with PBS containing 0.1% Tween 20, the sections were blocked in 5% BSA for 1 h and incubated overnight at 4°C with rabbit anti-cleaved-caspase-3 primary antibody (Servicebio) diluted in PBS with 0.1% Tween 20. After washing, the sections were incubated with secondary antibody (horseradish peroxidase-labeled sheep anti-rabbit IgG, Beijing ZSGB-BIO Co., Ltd.) for 45 min. Finally, the slides were developed using the 3,3’-diaminobenzidine (DAB), and hematoxylin was used for counterstaining. All immunohistochemical sections were scanned with a Leica Apero CS2 slide scanning system.

### In vivo propidium iodide (PI) staining assay of placental pyroptosis

Mock- or ZIKV-infected mice were administered PI (1.5 mg/kg) via the intravenous route at E16.5 before subjecting them to sacrifice. About 15 min later, mice were euthanized, and the placentas were collected and embedded in the O.C.T compound (SAKURA, 4583). After slicing, the slides were stained with Hoechst and scanned on EVOS FL auto (Thermo Fisher Scientific).

### In vitro RNA transcription and transfection

The DNA fragments corresponding to the 5’ UTR and 3’ UTR of ZIKV genome were cloned using the cDNAs from viral stocks. The 5’ and 3’ UTR fragments were amplified by PCR using the following primers: 5’ UTR forward primer 5’-CGGGGTACCAGTTGTTGATCTGTGTGAATCAGA-3’, 5’ UTR reverse primer 5’-CCGCTCGAGGACCAGAAACTCTCGTTTCCAAA-3’, 3’ UTR forward primer 5’-CGGGGTACCGCACCAATCTTAGTGTTGTCAGGCC-3’, and 3’ UTR reverse primer 5’-CCGCTCGAGAGACCCATGGATTTCCCCAC-3’. Amplified fragments were digested with KpnI and XhoI and cloned into the pre-cut pcDNA4/myc-His A vector (Invitrogen) by using the DNA ligation Kit (Takara). Plasmid DNA was then linearized with the restriction enzyme XhoI, and used as a template for T7 in vitro transcription using the mMESSAGE mMACHINE T7 Transcription Kit (Ambion). The RNA was precipitated with lithium chloride and quantified by spectrophotometry. Transfections were performed using the Lipofectamine 2000 (Invitrogen) according to the manufacturer’s instructions

### RNA extraction and quantitative real-time PCR

Total RNA in treated cells was extracted using TRIzol Reagent (Invitrogen), and 1 µg RNA was used to synthesize cDNA using a first-strand cDNA synthesis kit (TOYOBO). Quantitative real-time PCR was performed using a 7500 Real-Time PCR System (Applied Biosystems) and SYBR Green PCR Master Mix (TOYOBO). Data were normalized to the level of β-actin expression in each sample. The primer pairs used were as follows: β-actin: forward 5′-AGCGGGAAATCGTGCGTGAC-3′, reverse 5′-GGAAGGAAGGCTGGAA GAGTG-3′; TNF-α: forward 5′-CCTCTCTAATCAGCCCTCTG-3′, reverse 5′-GAGGACCTGGGAGTAGATGAG-3′.

### Statistical analysis

All data were analyzed with the GraphPad Prism software. For viral burden analysis, the log titers and levels of viral RNA were analyzed by nonparametric Mann-Whitney tests. Contingency data were analyzed by Fisher’s exact test using numbers in each group. The statistical analysis of differences between two groups was performed using two-tailed Student’s t-test. For all statistical significance indications in this manuscript, ****, p < 0.0001; ***, p < 0.001; **, p < 0.01; *, p < 0.05 and ns, no significance.

## Acknowledgments

We acknowledge F. Shao (National Institute of Biological Sciences, Beijing, China) for providing Gsdme^−/−^ mice.

## Author contributions

J.Y. and S.C. conceived and supervised the experiments. Z.Z., Q.L., M.Y., W.Z. and Z.C. performed the experiments. Z.Z. and J.Y. planned experiments and analyzed results. Z.Z., J.Y. and U.A. wrote the manuscript. J.Y. and S.C. checked and finalized the manuscript. All authors read and approved the final manuscript.

---

**Fig. S1.**
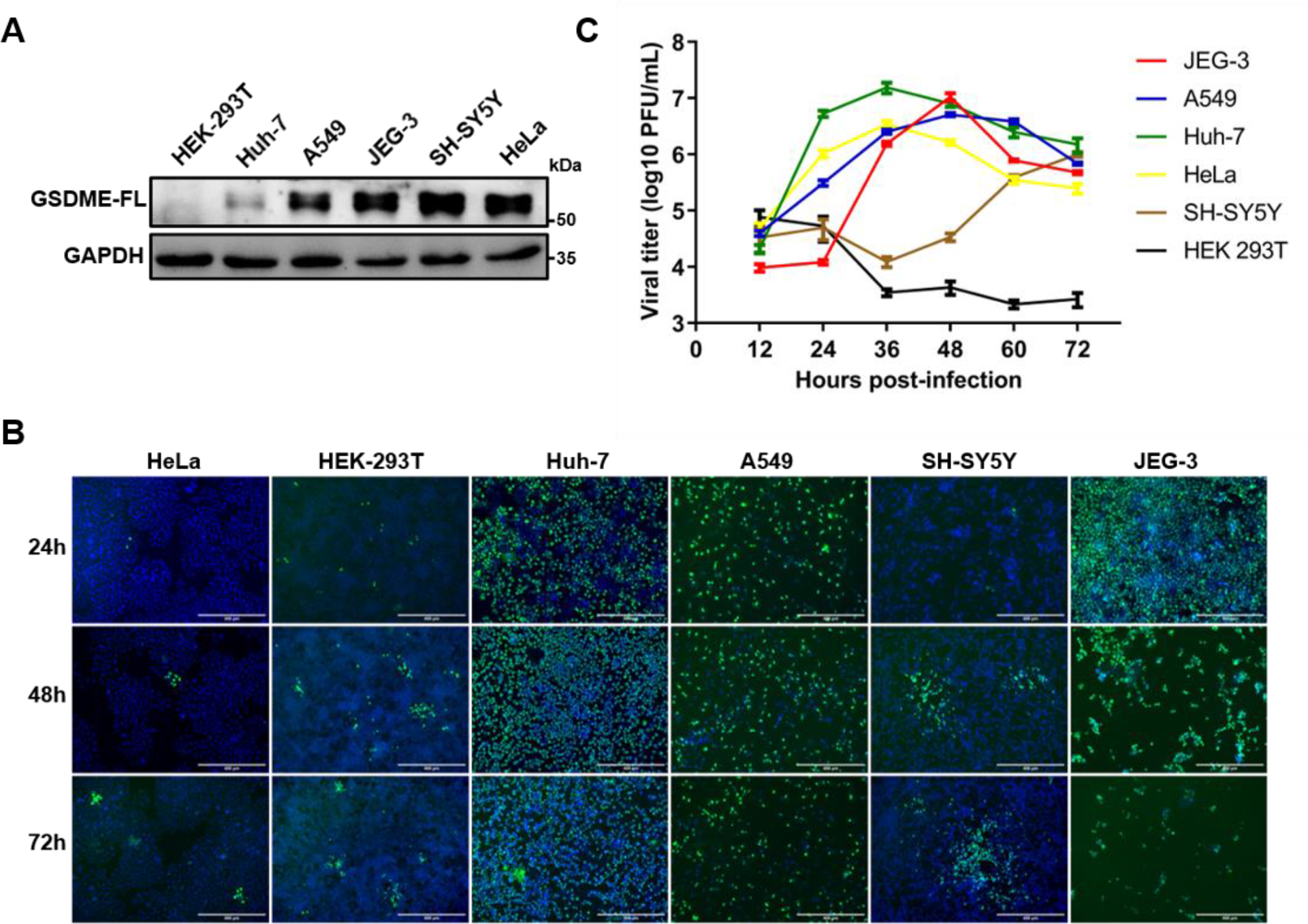
The GSDME abundance and the susceptibility to ZIKV infection of relevant cell lines. **(A)** Immunoblot analyses of GSDME-FL in indicated cell lines. **(B to C)** Relevant cells were infected with ZIKV at a MOI of 0.1. At indicated time post-infection, cells were subjected to immunofluorescence analysis (B). Scale bar, 400 μm. For plaque assay, the supernatants of ZIKV-infected cells were harvested at 12-72 h post-infection, and the titration was performed on Vero cells (C).

**Fig. S2.**
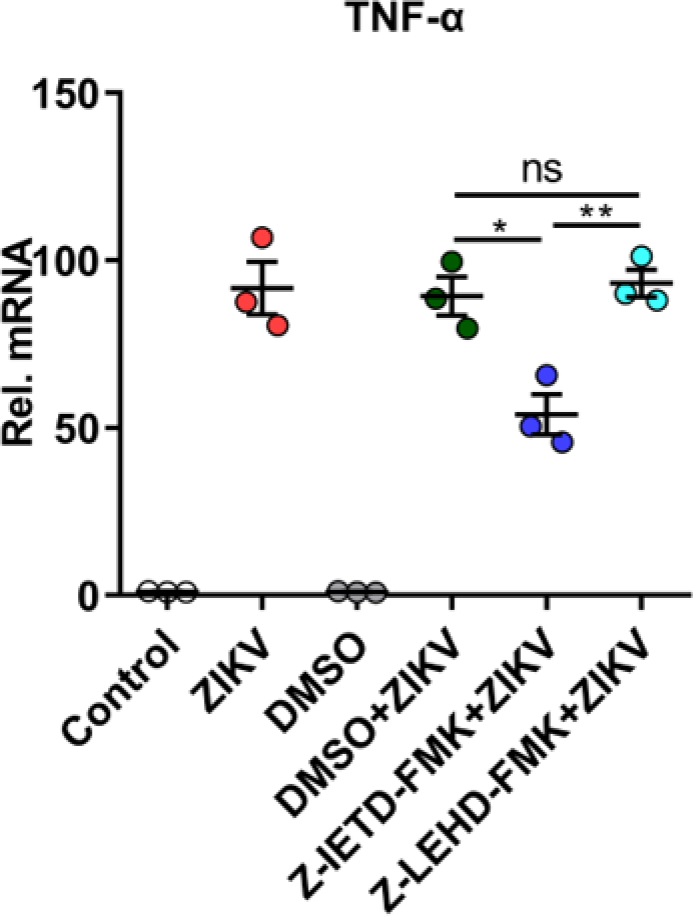
ZIKV infection significantly enhances the expression of TNF-α in infected JEG-3 cells. JEG-3 cells were infected with ZIKV at a MOI of 1, and were subsequently incubated with either 25 μM Z-IETD-FMK or 25 μM Z-LEHD-FMK for 24 h. Then the cells were harvested for RNA extraction and subjected to detect mRNA level of TNF-α by qRT-PCR. Data were normalized to the level of β-actin expression in each sample. All data are presented as the mean ± SEM of at least three independent experiments. **, p< 0.5; **, p < 0.01; ns, no significance.

**Fig. S3.**
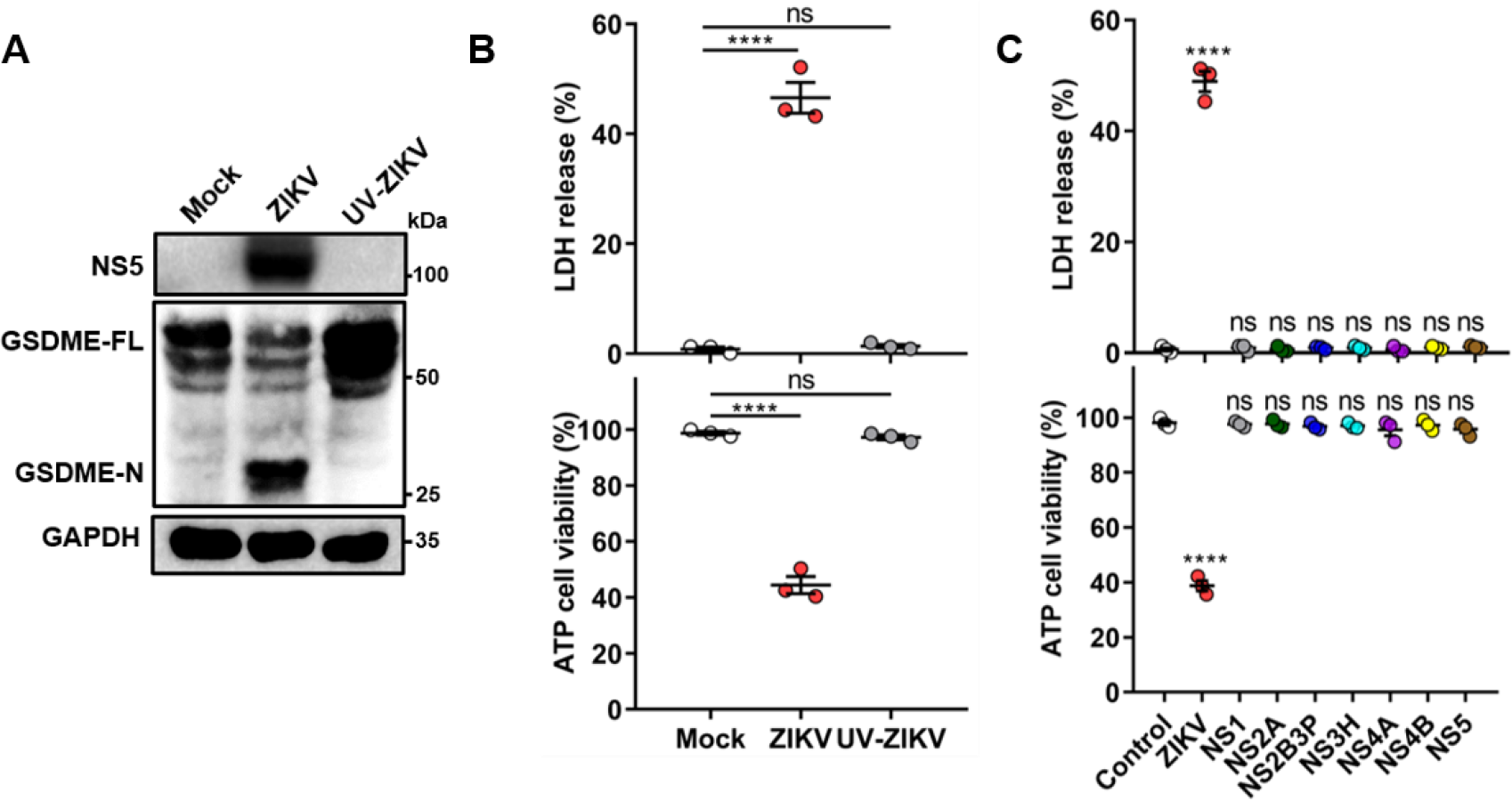
The structural and non-structural proteins of ZIKV are not capable of inducing pyroptosis. **(A to B)** JEG-3 cells were infected with ZIKV or UV-inactivated ZIKV at a MOI of 1. At 24 h post-infection, cells were subjected to western blot (A) and cytotoxicity analyses (B). **(C)** JEG-3 cells were seeded in 6-well plate and transfected with 2 μg indicated plasmids. At 24 h post-transfection, cells were subjected to cytotoxicity analyses. Unpaired t test versus Control. All data are presented as the mean ± SEM of at least three independent experiments. ****, p < 0.0001; ns, no significance.

**Fig. S4.**
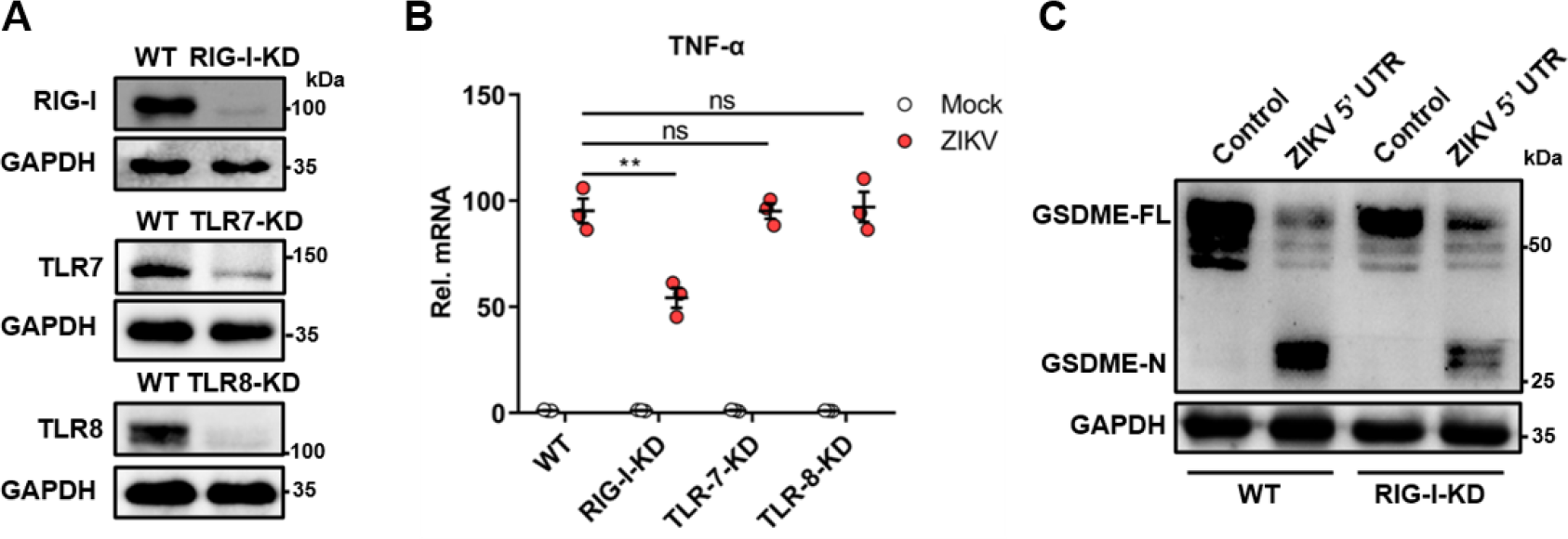
The KD of RIG-I suppresses the ZIKV or ZIKV 5’ UTR-induced activation of pyroptosis. **(A)** Verification of RIG-I, TLR7and TLR8 knockdown efficiency by immunoblotting. **(B)** JEG-3 WT or RIG-I KD cells were seeded in 6-well plate and transfected with 1 μg ZIKV 5’ UTR. At 24 h post-transfection, the cells were harvested for western blot analyses. **(C)** Relevant cells were infected with ZIKV at a MOI of 1. At 24 h post-infection, the cells were harvested for RNA extraction and subjected to detect mRNA level of TNF-α by qRT-PCR. Data were normalized to the level of β-actin expression in each sample. All data are presented as the mean ± SEM of at least three independent experiments. **, p < 0.01; ns, no significance.

**Fig. S5.**
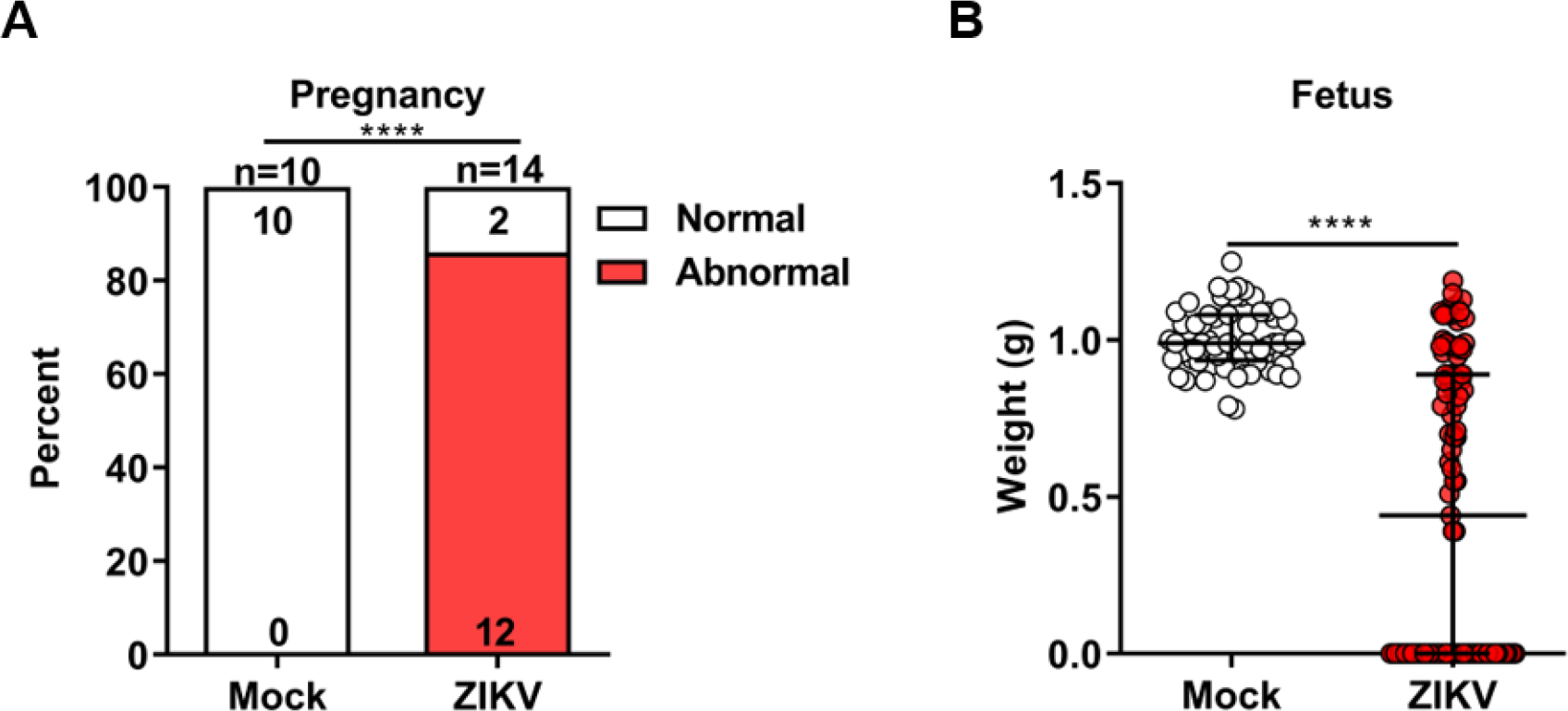
Impact of ZIKV on dams and fetuses at E16.5. Pregnant dams were infected with ZIKV or Vero cell culture supernatant (Mock) as described in Fig. 5A. **(A)** The percentage of dams showing abnormal pregnancy (at least one fetus showed morphological abnormality or suffered demise/resorption). Numbers on bars indicate normal pregnancy (top) or abnormal pregnancy (bottom). Significance was determined by Fisher’s exact test. ****, p<0.0001. **(B)** The weight of fetuses. Data shown are median with interquartile range, and Mann-Whitney test was used to calculate significance. ****, p < 0.0001. Data are pooled from 3-5 independent experiments.

**Fig. S6.**
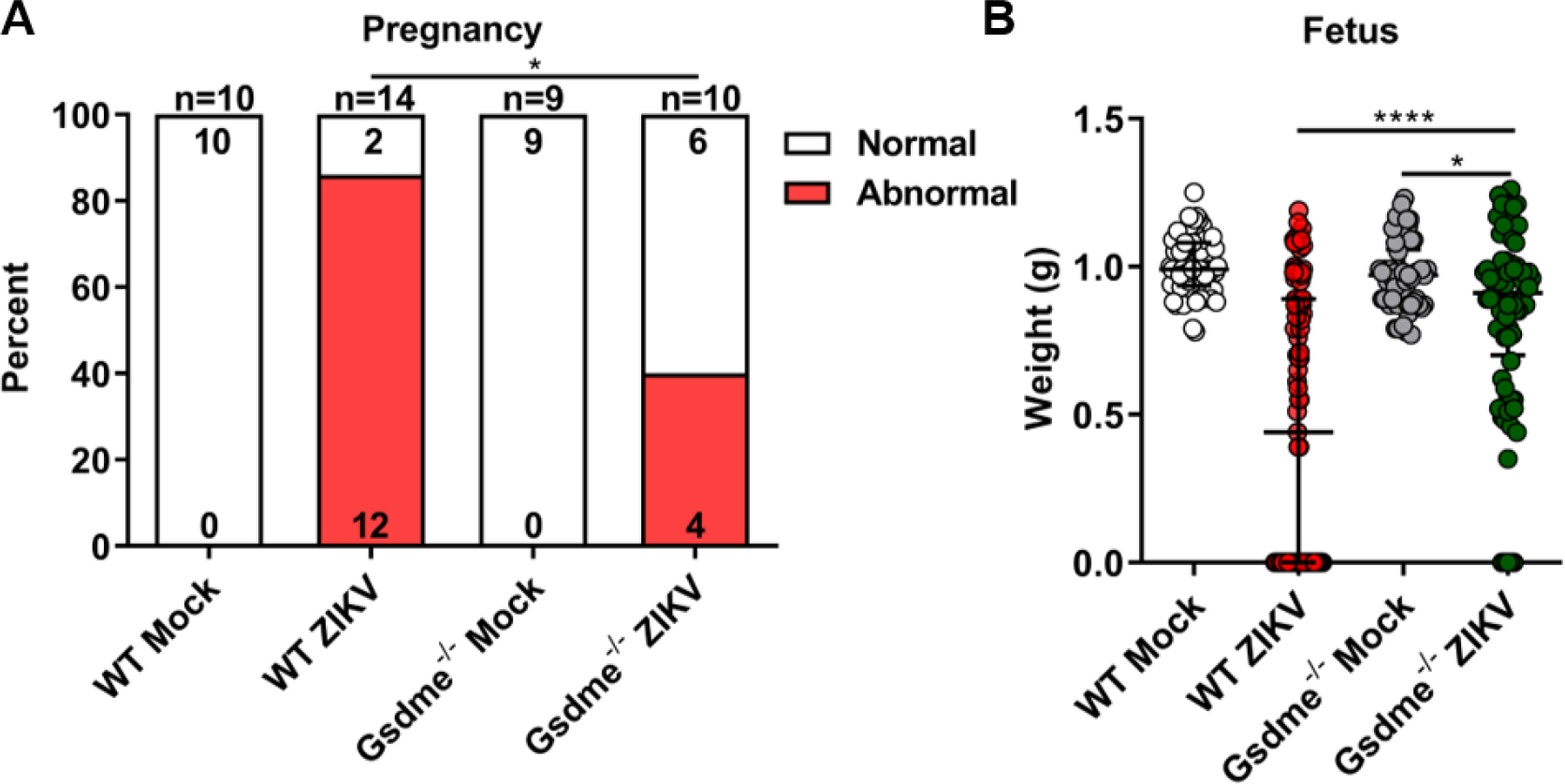
GSDME deletion restrains ZIKV-induced abnormal pregnancy and adverse fetal outcomes. Pregnant WT and Gsdme^-/-^ dams were infected with ZIKV or Vero cell culture supernatant (Mock) as described in Fig. 5A. **(A)** The percentage of dams showing abnormal pregnancy (at least one fetus showed morphological abnormality or suffered demise/resorption). Numbers on bars indicate normal pregnancy (top) or abnormal pregnancy (bottom). Significance was determined by Fisher’s exact test. *, p<0.05. **(B)** The weight of fetuses. Data shown are median with interquartile range, and Mann-Whitney test was used to calculate significance. *, p<0.05; ****, p < 0.0001. Data are pooled from 3-5 independent experiments.

